# Characterization of Chromatin-Associated Circular RNAs (cacRNAs)

**DOI:** 10.64898/2026.07.20.739476

**Authors:** Susovan Sadhukhan, Komal Kumari, Pranita Rout, Amaresh Chandra Panda

## Abstract

**Highlights:**

- Identified hundreds of potential chromatin-associated circRNAs in HEK293 cells, H9, and HeLa cells
- The first report suggesting global interaction of circular RNAs with chromatin
- Chromatin-associated circular RNAs interact with various RBPs involved in RNA splicing or processing

Circular RNAs (circRNAs) have emerged as novel regulators of gene expression by interacting with various proteins and RNAs in a spatiotemporal manner. CircRNAs localized in the cytoplasm regulate mRNA translation or stability by binding to microRNAs and RNA-binding proteins (RBPs), while circRNAs in the nucleus regulate transcription and pre-mRNA splicing by associating with transcription factors and splicing factors. In this study, we sought to explore the interaction between circRNAs and chromatin. Analyzing published RNA-seq data from chromatin fractions identified hundreds of chromatin-associated circRNAs (cacRNAs) in various human cells. We validated the enrichment of a subset of circRNAs in the chromatin fraction and established the direct interaction of *circDYNC1H1* and *circKIF2C* with chromatin in HEK293T cells. Furthermore, cacRNAs were found to interact with RBPs. Together, our research demonstrates the global association of hundreds of circRNAs with chromatin and expands our understanding of novel functional aspects of the circRNAs.

Graphical abstract

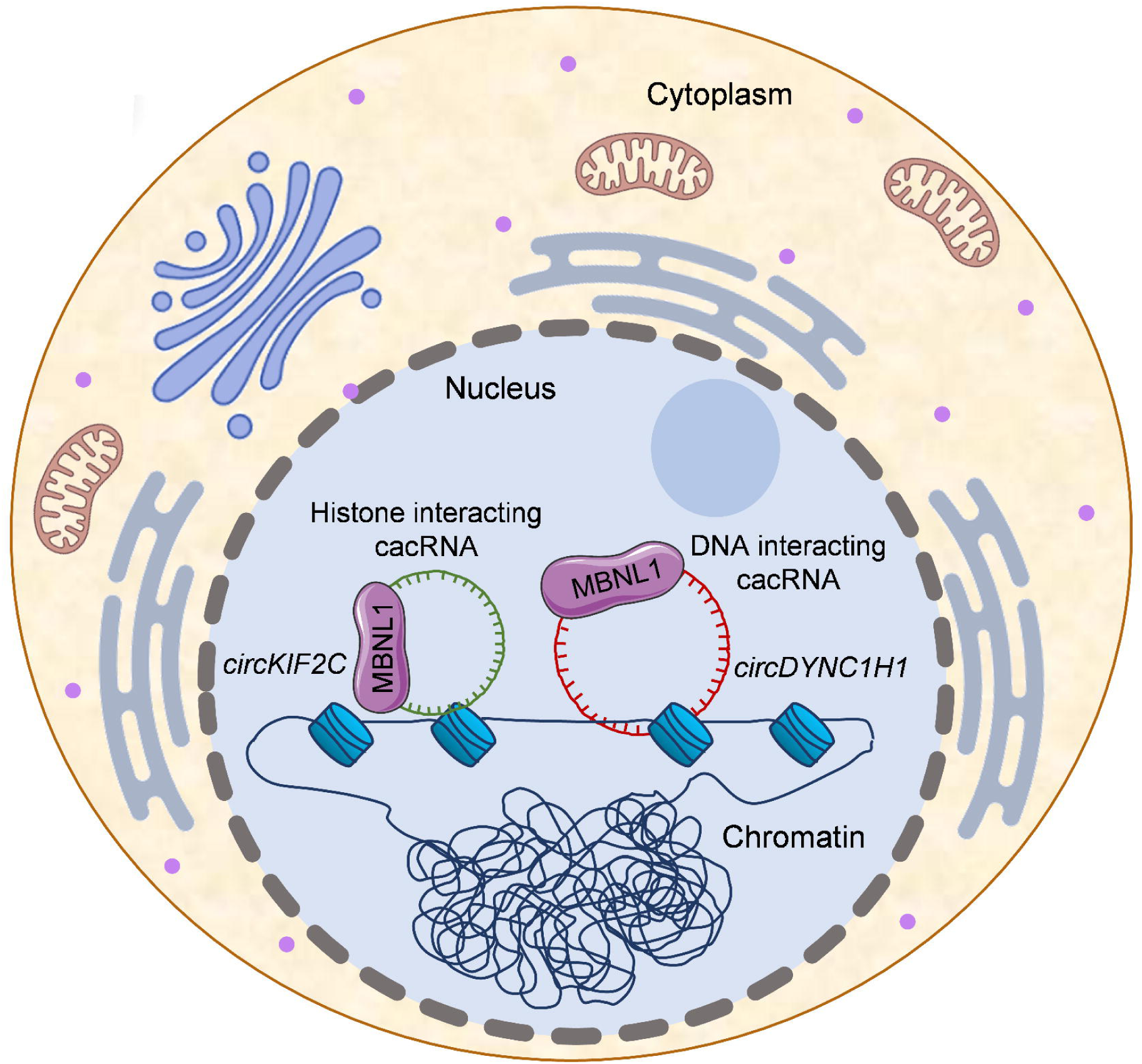

## INTRODUCTION

CircRNAs are a covalently closed family of RNA molecules without any 5’ cap and poly-A tails and are ubiquitously expressed across all eukaryotes [1, 2]. CircRNAs are generated by the backsplicing process from pre-mRNAs and are resistant to exonucleases due to their closed-loop structure [3, 4]. Over the past decade, numerous studies have established that circRNAs are a critical regulator of cellular functions [2].

After their biogenesis through backsplicing in the nucleus, circRNAs get localized to different cellular compartments and interact with various biomolecules like micro (mi)RNAs, RNA-binding proteins (RBPs), and mRNAs to regulate gene expression [5–9]. Several recent studies have also reported the translation of circRNAs into functional proteins that regulate cellular physiology [10, 11]. A few recent reports suggested that various regulatory RNA molecules can regulate gene expression by interacting with chromatin [12]. Global RNA interaction with DNA sequencing (GRID-seq) and chromatin-associated RNA sequencing (ChAR-seq) discovered several chromatin-associated RNAs, including nascent transcribing coding and noncoding (nc)RNAs, snRNAs, snoRNAs, rRNAs, and lncRNAs [13, 14]. Furthermore, Profiling Interacting RNAs on Chromatin followed by deep sequencing (PIRCh-seq) suggested the association of various ncRNAs with different modified histones [15]. Cetin *et al.* have developed a method to identify RNAs that form DNA-RNA triplexes across the genome [16]. Another study performed biochemical fractionation of the nuclear compartment and identified a distinct subclass of lncRNAs that interact with RNA pol II and are tethered to chromatin near active genes [17]. These studies established the critical role of chromatin-associated RNAs in chromatin architecture and the regulation of gene expression, adding a new dimension to the genetic interactome.

Only a few studies have reported associations between circRNA molecules and cis-DNA sequences and/or RNA pol II in regulating the transcription of their host genes [18, 19]. For example, *circSMARCA5* downregulates *SMARCA5* transcription by pausing RNA pol II elongation, whereas *circPAIP2* and *ci-ankrd52* interact with RNA pol II and upregulate transcription of their parental genes [20–22]. However, the global interaction between circRNAs and chromatin remains unclear. In our study, we analyzed previously published RNA-sequencing data from chromatin fractions of HEK293 cells, histone-associated RNAs from H9 cells, and DNA-RNA triplex-forming RNAs from HeLa cells, and found that hundreds of circRNAs are enriched in chromatin. A subset of chromatin-associated circRNAs (cacRNAs) were validated in HEK293T cells, and biochemical and computational analyses assessed their association with chromatin. Furthermore, the analysis of cacRNA interactions with RBPs indicated their association with various proteins involved in chromatin-level functions, such as RNA processing and RNA export. Overall, our data provide evidence of global circRNA-chromatin association and its potential to regulate gene expression.

## MATERIALS AND METHODS

### Cell culture and subcellular fractionations

HEK293T cells were cultured in Dulbecco’s Modified Eagle Medium (DMEM) supplemented with 10% Fetal Bovine Serum (FBS) and 1% penicillin–streptomycin, kept in an incubator at 37°C and 5% CO2. For Actinomyocin D (ActD) treatment, cells were treated for 2 h before proceeding to further experiments. For subcellular fractionation, cells were pelleted down and lysed for 5 minutes on ice using 200 µL cytoplasmic lysis buffer (0.15% NP-40, 150 mM NaCl, 1x protease inhibitor, and 20 U of RNase inhibitor). The cell lysate was layered onto 500 µL of sucrose buffer (10 mM Tris-Cl, pH 7.0, 150 mM NaCl, and 25% w/v sucrose), then centrifuged at 16,000g for 10 minutes at 4 °C. The cytoplasmic supernatant was taken out in a separate tube, and the nuclear pellet was washed with 800 μL wash buffer (0.1% v/v Triton X-100 and 1 mM EDTA dissolved in 1x PBS) by centrifugation at 1,150 *g* for 1 min at 4 ^°^C. Then the nuclear pellet was resuspended in 200 µL of glycerol buffer (20 mM Tris-cl pH8.0, 75 mM NaCl, 0.5 mM EDTA, 50% v/v glycerol, 0.85 mM DTT) and the nuclei was lysed with 200 µL nuclei lysis buffer (1% v/v NP-40, 20 mM HEPES pH 7.5, 300 mM NaCl, 1 M urea, 0.2 mM EDTA, 1 mM DTT). The sample was centrifuged at 18,500 *g* for 2 minutes at 4 ^°^C, followed by the collection of the nuclear supernatant and chromatin pellet for RNA isolation [23].

### RNA-sequencing Analysis

SRA files for the chromatin fractions of HEK293 cells (GSE66478), PIRCh-seq (GSE119006), and DNA-RNA triplex (GSE120850) were downloaded from the NCBI database and converted to FASTQ, followed by quality assessment using FastQC. The clean reads were then aligned to the human genome (hg38) using the STAR aligner. This alignment information was parsed and annotated through CIRCexplorer2 (v2.3.8) for identification of circRNAs, and all circRNA annotation files were compiled using rbind [24]. We checked the abundance of circRNAs in different fractions of HEK293 cells using the TPM (transcripts per million) method and calculated the z-score using edgeR (**Supplementary Table S1**) [25]. The list of cacRNAs expressed in H9 and HeLa cells are listed in **Supplementary Table S2**.

### RNA isolation, RT-PCR, and Sanger sequencing

The total RNA was isolated using MagSure All RNA isolation kit (RNA Biotech). The RNAs were then converted to cDNA using Maxima reverse transcriptase following the manufacturer’s protocol (Thermo Fisher Scientific). PowerUp SYBR Green PCR Master Mix and divergent primers (**Supplementary Table S3**) were used to amplify the specific amplicon under the following PCR conditions: 95°C for 2 min, followed by 40 cycles of 95°C for 5 s and 60°C for 20 s. The PCR products were then checked on agarose gel, and purified PCR products were subjected to Sanger sequencing using one of the divergent primers to identify circRNA back-splicing junction (BSJ) sequences [26].

### RNase R treatment and quantitative PCR

2 ug of the total RNAs were treated with 0.25 µL RNase R enzyme for 30 minutes at 37 °C. The treated RNA samples were used to perform RT-qPCR for the linear and circRNAs. For quantitative analysis, PowerUp SYBR Green Master Mix (Thermo Fisher Scientific) was used to amplify the circRNA BSJ sequences using divergent primer pairs under real-time PCR conditions of 95°C for 2 min, followed by 40 cycles of 95°C for 5 s and 60°C for 20 s. Relative change was calculated using the ΔΔCT method [27].

### Western Blot

For performing western blot from the chromatin fraction, the chromatin samples were subjected to sonication using Bioruptor sonication device (Diagenode). The lysates were loaded onto an SDS-PAGE gel and transferred to a nitrocellulose membrane. The membrane was incubated overnight at 4 ^°^C with specific primary antibodies for GAPDH (CST #2118), Histone H3 (CST #14269), GFP (Abclonal #AE078), and MBNL1 (CST #94633), then with appropriate HRP-conjugated secondary antibodies, and bands were visualized using a chemiluminescent kit (Promega).

### DNA-RNA hybrid pulldown

For DNA-RNA hybrid pulldown, the chromatin pellet from HEK293T cells was resuspended in PEB buffer (20 mM Tris-Cl pH 7.5, 100 mM KCl, 5 mM MgCl2, and 0.5% NP-40) and sonicated using a Bioruptor sonication device for 10 pulses of 30 seconds each. The Protein G beads (Thermo Scientific) were washed, resuspended in NT2 buffer (50mM pH 7.4 Tris-Cl, 150mM NaCl, 1mM MgCl_2,_ and 0.05% NP-40), and incubated with control IgG (CST #5415) or S9.6 antibodies (Sigma #MABE1095) for 2 hours at room temperature [28]. After washing with NT2 buffer, the bead-antibody complexes were further incubated with sonicated chromatin samples for 2 hours at 4 °C. After washing, the RNAs from the pulldown samples were isolated for further analysis.

### Protein and DNA interaction analysis

Ribonucleoprotein Immunoprecipitation (RIP) Analysis was performed in HEK293T cells as described previously with little modifications [29]. The chromatin pellet was subjected to sonication in PEB buffer. The protein G beads (Thermo Scientific) were incubated with the specific antibodies (2 µg) for 2 hours at room temperature, then incubated with the chromatin lysate for 2 hours at 4 °C. After that, the beads were washed 3 times with NT2 buffer, followed by western blot and qPCR analyses. For the genetic interaction of circRNAs, we performed local nucleotide BLAST analysis (BLASTn) of the circRNA sequences against the database generated from GRCh38.primary_assembly.genome.fa as the reference **(Supplementary Table S4)**. Triplex formation analysis was performed using the 3plex Web server, with the circRNA sequences of *circDYNC1H1* and MANE (len: 19205) as the DNA target site [30]. To predict protein interactions with cacRNAs, the list of RBPs was extracted from the CIRCpedia v3 **(Supplementary Table S5)** [31].

### Gene Ontology (GO), and Kyoto Encyclopedia of Genes and Genomes (KEGG) Pathway analysis

The predicted proteins were used as input to the DAVID database to perform GO enrichment and KEGG pathway analyses [32]. The enrichment score for all GO terms is calculated as the -log10 of the p-value **(Supplementary Table S6)** [33, 34].

### Visualization and statistical analysis

The graphs were created using various tools, including Microsoft Excel, Microsoft PowerPoint, GraphPad Prism, and RStudio. All experiments had at least 3 biological replicates, and error bars represent the mean ± standard error of the mean (SEM). A Student’s t-test was performed to calculate the p-value, and p-values < 0.05 were considered significant.

## RESULTS

### CircRNA analysis reveals thousands of chromatin-associated circRNAs (cacRNAs)

As chromatin is mainly composed of histones and DNA, the cacRNAs might interact directly with histones, chromatin-localized proteins, or DNA. To identify cacRNAs, we selected three previously published RNA-seq data sets of RNAs isolated from the total chromatin fraction of HEK293 cells (GSE66478), RNAs associated with different modified histones (PIRCh-seq) of H9 cells (GSE119006), and RNAs from the DNA-RNA triplex of HeLa cells (GSE120850) **(Figure 1A)** [15–17]. The analysis of these RNA-seq samples using CIRCexplorer2 identified 1794 circRNAs in the HEK293 chromatin sample, 385 circRNAs with histones in the PIRCh-seq data of H9 cells, and 1613 circRNAs in DNA-RNA triplexes of HeLa cells **(Figure 1A)**. As circRNAs show tissue-specific expression patterns, only a small fraction of cacRNAs were common in these three types of samples [35]. There are 46 cacRNAs common between HEK293-chromatin and H9-histones, 95 common cacRNAs between HEK293-chromatin and DNA-RNA triplexes of HeLa cells, and 21 common cacRNAs between H9-histones and HeLa-DNA-RNA triplexes, indicating conserved and functionally relevant sets of cacRNAs interacting with histones and DNA **(Figure 1B-D)**. The circos plots show the origins of cacRNAs from different chromosomes across all three datasets **(Supplementary Figure S1A-C)**.

**Figure 1:**
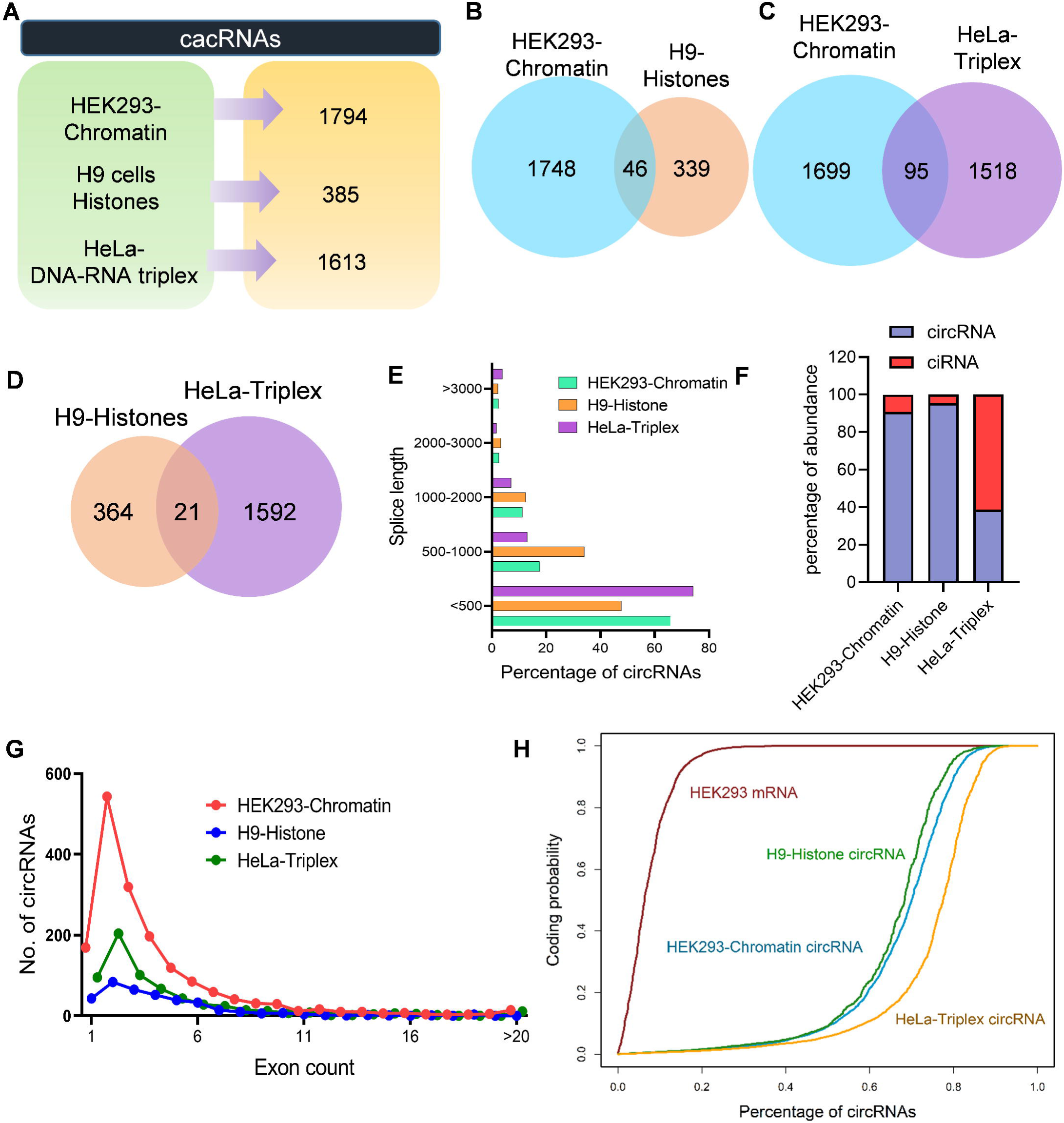
Identification of cacRNAs from RNA-seq data. **A.** Schematic showing the numbers of cacRNAs identified in the RNA-seq data of HEK293-chromatin, H9-Histone PIRCh-seq, and HeLa-DNA-RNA triplex samples in the mentioned cell lines. **B-D.** The Venn diagram showing the common circRNAs between HEK293-chromatin circRNAs and H9 histones circRNAs (B), HEK293-chromatin circRNAs and HeLa-Triplex circRNAs (C), and H9-histones circRNAs and HeLa-Triplex circRNAs (D). **E.** The distribution of cacRNA lengths across all three RNA-seq samples. **F.** The fractions of exonic and intronic cacRNAs of the three RNA-seq data. **G.** The distribution of exons in the exonic cacRNAs from the three RNA-seq datasets. **H.** Coding probability of the HEK293-mRNAs, HEK293-Chromatin circRNAs, H9-Histone circRNAs, and HeLa-Triplex circRNAs.

We compared our annotated cacRNAs with the circRNAs from circBase, circAtlas, and CIRCpedia v3 database and found that 654 DNA-RNA triplex-forming circRNAs in HeLa, 34 chromatin-associated circRNAs in HEK293, and 16 histone-associated circRNAs in H9 cells were novel circRNAs **(Supplementary Figure S1D-F)**. We found a similar length distribution across all cacRNA types, and most cacRNAs are shorter than 1000 nucleotides **(Figure 1E)**. Among these cacRNAs, chromatin- and histone-associated circRNAs have higher exonic circRNAs, but, surprisingly, triplex-forming circRNAs have higher intronic circRNAs **(Figure 1F)**. However, the analysis of exonic cacRNAs revealed that in all three cases, most cacRNAs have fewer than 10 exons **(Figure 1G)**. Some recent articles claim that circRNAs can be translated into proteins [10]. As the cacRNAs reside within the nucleus, they must have lower coding probabilities. As expected, CPAT analysis revealed that HEK293 mRNAs have very high coding probabilities, whereas cacRNAs have very low coding probabilities **(Figure 1H)**.

### Validation of the cacRNAs in HEK293T cells

To further validate the cacRNAs, we focused on the chromatin RNA-seq data, as these circRNA pools contain both histone- and DNA-interacting circRNAs. In the previous study, RNA-seq was performed on soluble nuclear extract (SNE), chromatin pellet extract (CPE), and DRB-treated chromatin pellet extract (DRB) [17]. After analysis of circRNAs, we found differentially expressed circRNAs between the SNE and CPE samples **(Figure 2A)**. Individually, there are 1514 circRNAs in SNE, 944 circRNAs in CPE, and 966 circRNAs in DRB samples, and 62 circRNAs were common between all three samples **(Figure 2B)**. These common circRNAs were showing significant enrichment in the chromatin fraction **(Figure 2C)**. We also found strong negative correlations between SNE and CPE, and between SNE and DRB circRNAs, indicating an overall lower abundance of circRNAs in chromatin. As expected, there was also a strong negative correlation between CPE vs DRB circRNAs, as the DRB inhibits the transcription and lowers the expression of circRNAs in the chromatin fraction **(Figure 2D)**. The circRNAs from all these three sources show similar trends in splice length, exon counts, and the ratio of the exonic and intronic circRNAs **(Supplementary Figure S2A-I)**.

**Figure 2:**
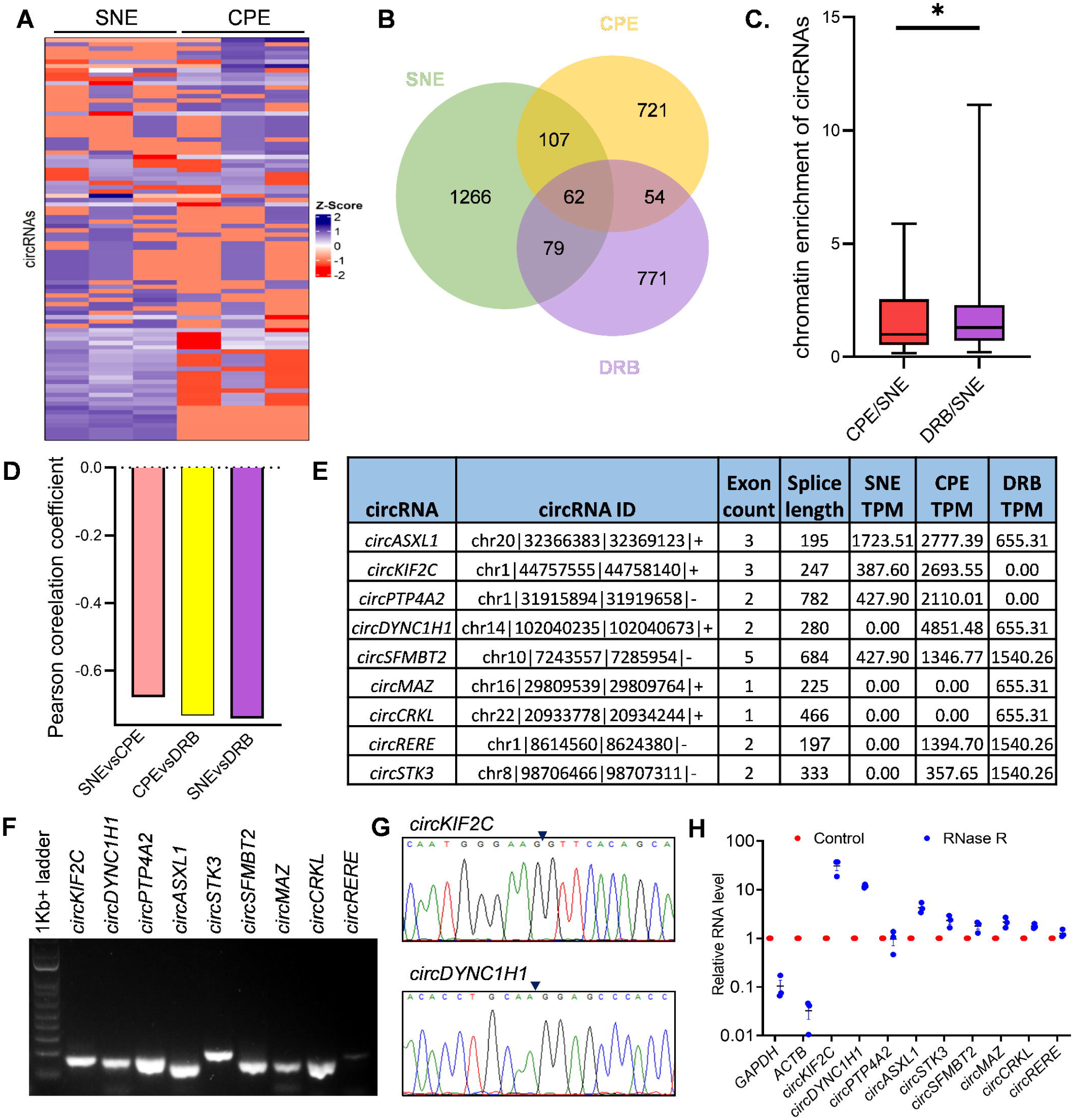
Validation of cacRNAs in HEK293T cells. **A.** Heatmap showing the differential abundance of circRNAs between SNE and CPE samples. **B.** Venn diagram showing the common circRNAs between SNE, CPE, and DRB samples. **C.** Box plot showing the chromatin enrichment of common circRNAs calculated from the TPM of the individual circRNAs. **D.** Pearson correlation coefficient between SNE vs CPE, CPE vs DRB, and SNE vs DRB circRNAs. **E.** Selected circRNAs for further analysis. **F.** RT-PCR product of the circRNAs using BSJ-specific divergent primers. **G.** Sanger sequencing results to confirm the BSJ sequence of the circRNAs (the arrowhead denotes the BSJ point). **H.** RT-qPCR analysis showing the level of the linear RNAs and circRNAs after treatment of RNase R in total RNAs. Data represent the ± SEM of six biological replicates.

To validate chromatin-associated circRNAs, we selected a subset of 9 circRNAs based on the following criteria: higher abundance either in the CPE or DRB samples compared to SNE samples, shorter splice length, and fewer exon counts **(Figure 2E)**. We amplified BSJs of the circRNAs and performed PCR to validate their expression in the HEK293T cells **(Figure 2F)**. Sanger sequencing further confirms amplification of the backsplicing junction of the circRNAs (**Figure 2G, Supplementary Figure S3)**. The RNase R exonuclease degraded linear RNAs such as *GAPDH* and *ACTB* mRNA, while the circRNAs showed resistance to RNase R digestion, confirming their closed-loop structure **(Figure 2H)**.

### Chromatin fraction contains cacRNAs

We fractionated the cells to get cytoplasmic, nuclear, and chromatin fractions, followed by RNA isolation **(Figure 3A, B)**. The quality of the fractions was confirmed by the cytoplasmic enrichment of GAPDH protein and enrichment of Histone H3 protein in chromatin fractions **(Figure 3C)**. The subcellular separation was further confirmed by RT-qPCR validation of specific enrichment of *GAPDH* mRNA in cytoplasm, *MALAT1* lncRNA in nuclear supernatant, and *U6* in the chromatin fractions **(Figure 3D)**. Within these fractions, we checked the comparative abundance of the selected circRNAs, and the results revealed that although the circRNAs are present in the cytoplasm and nuclear lysates, a significant percentage of some of the circRNAs remain within the chromatin fraction **(Figure 3E, Supplementary** F**igure S4A)**.

**Figure 3:**
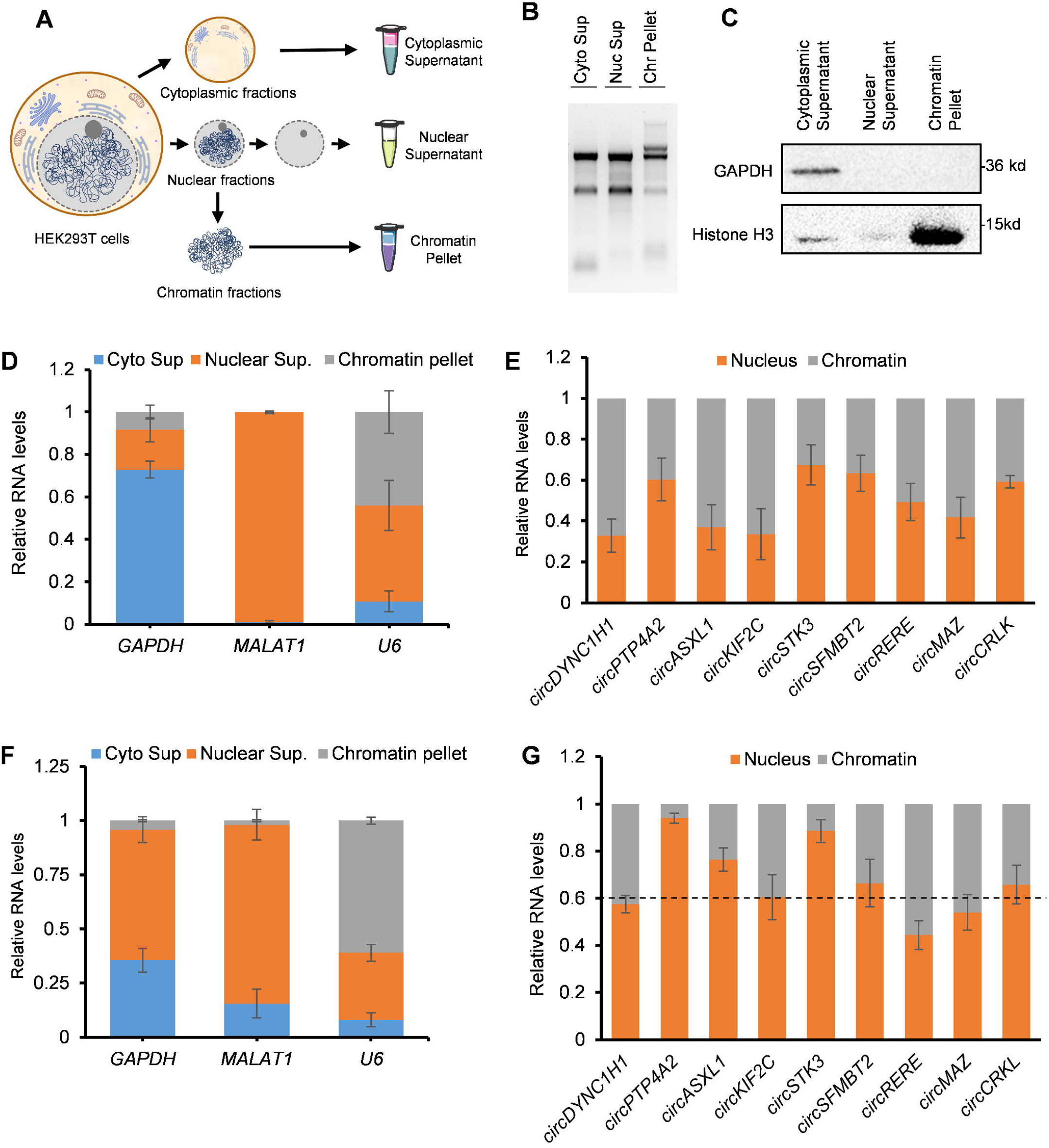
Biochemical separation of HEK293T cells showing abundance of the circRNAs with chromatin. **A.** Schematic showing cell fractionation followed by RNA isolation. **B.** Agarose gel electrophoresis of the total RNAs isolated from the three cell fractions-cytoplasmic supernatant, nuclear supernatant, and chromatin pellet. **C.** Western blot of specific cell fractionation markers to ensure proper cell fractionation. **D.** RT-qPCR analysis of the cell fractionation RNA markers. **E.** RT-qPCR analysis showing the level of circRNAs in nuclear supernatant and chromatin pellet fractions. **F.** RT-qPCR analysis of the cell fractionation RNA markers in Actinomycin D-treated cells. **G.** RT-qPCR analysis showing the level of circRNAs in nuclear supernatant and chromatin pellet fractions in Actinomycin D-treated cells. Data in panel D-G represent the average ± SEM of four biological replicates.

As circRNA biogenesis is a co-transcriptional process, assessing circRNA association with chromatin via cellular fractionation may yield false-positive results [36]. To investigate the genuine interaction of circRNA with chromatin DNA, we treated the cells with Actinomycin D to inhibit transcription, followed by subcellular fractionation, RNA isolation, and verification of subcellular RNA markers to confirm the quality of the fractionation (**Figure 3F, Supplementary Figure S4B)**. As expected, most of the circRNAs get depleted from the chromatin fractions in the absence of transcription, while *circDYNC1H1, circKIF2C, circRERE,* and *circMAZ* were found to remain associated with the chromatin **(Figure 3G, Supplementary Figure S4C).**

### Interaction of cacRNAs with chromatin DNA and histone

As chromatin is primarily composed of histones and DNA, we performed pulldown assays to assess whether the abundant cacRNAs interact with histones or DNA **(Figure 4A)**. As the C-terminal tail of histone 3 (H3) is tri-methylated at the 4^th^ lysine residue (H3K4me3) in the promoter region to activate downstream gene transcription, we performed a pulldown of H3K4me3 to assess cacRNA association with actively transcribed promoter regions. Interestingly, pulldown of H3K4me3-modified histone from chromatin, followed by RT-qPCR analysis, revealed enrichment of *circKIF2C* and *circDYNC1H1* in the pulldown samples, indicating their interaction with histones **(Figure 4B, C)**. Furthermore, to assess the possibility of cacRNA interaction with chromatin DNA via R-loop formation, we performed DRIP-qPCR on the chromatin fraction using the S9.6 antibody [18, 37]. Interestingly, *circDYNC1H1* was found to be highly enriched in the S9.6 pulldown samples, suggesting that *circDYNC1H1* can form DNA-RNA hybrids, possibly forming the R-loop **(Figure 4D)**. We transfected HEK293T cells with the pEGFP-N2-2XNLS-RNaseH1 delta 1-27 (D210N) plasmid, which expresses a catalytic mutant RNase H in the nucleus [38]. After transfection, the protein is visible only in the nucleus and binds to DNA-RNA hybrids in chromatin **(Supplementary Figure 5)**. As expected, pulldown of RNaseH-GFP protein followed by RT-qPCR revealed the enrichment of *circDYNC1H1* RNaseH-GFP, further validating its direct association with DNA strands **(Figure 4E, F)**. Altogether, our results suggest that *circDYNC1H1* is a potent cacRNA that remains associated with both histone and chromatin DNA. The expression analysis of *circDYNC1H1* and *circKIF2C* in CIRCpedia v3 revealed that both are expressed in almost all human tissues **(Supplementary** f**igure 6A-B)**. In addition, these two circRNAs are well conserved in mammals, indicating their important role in higher organisms **(Supplementary** f**igure 6C-D)**.

**Figure 4:**
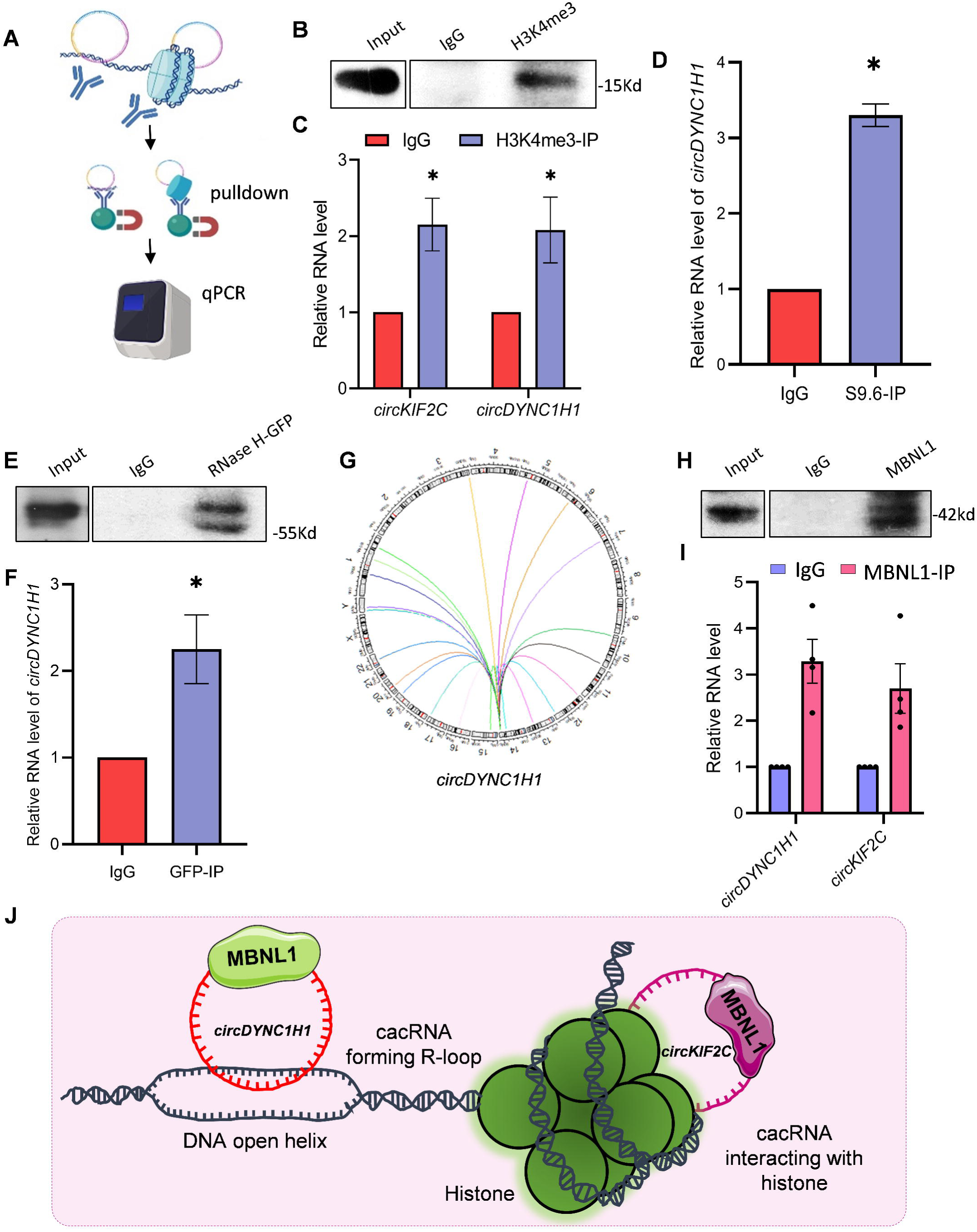
Direct interaction of *circKIF2C* and *circDYNC1H1* with chromatin. **A.** Schematic showing the pulldown of histone and DNA-RNA hybrids. **B.** Confirming H3K4me3 pulldown by western blot. **C.** H3K4me3 pulldown followed by qPCR. **D.** DRIP qPCR analysis by S9.6 antibody of the HEK293T cells. **E.** Pulldown of RNaseH-GFP protein using GFP antibody followed by western blot. **F.** Pulldown of RNaseH-GFP protein using GFP antibody followed by qPCR. **G.** Circos plot showing probable associated genomic regions of *circDYNC1H1*. **H.** MBNL1 pulldown followed by western blot. **I.** RT-qPCR analysis of circRNAs in MBNL1 pulldown samples. **J.** Proposed model on the association of cacRNAs with chromatin DNA and histones.

### Chromatin level functions of cacRNAs

As *circDYNC1H1* was found to be a DNA-interacting circRNA, we next sought to identify its likely interacting DNA regions. For that, we performed a global BLAST analysis, which predicted that *circDYNC1H1* can interact with 31 possible regions (**Figure 4G, Supplementary Table S4)**. Interestingly, *circDYNC1H1* is identified in triplex RNA-seq data we analyzed. Therefore, we wanted to check the possibility of triplex formation in the human genome. The 3plex web has predicted a strong triplex-forming domain in *circDYNC1H1*, spanning nucleotides 97 to 116 **(Supplementary Figure S7)** [30]. In addition, the BLAST analysis revealed three possible genomic regions complementary to *circDYNC1H1* that can form a DNA-RNA triplex, as predicted by the 3plex web server (**Supplementary Table S4**). Overall, all these data strongly suggest that *circDYNC1H1* can form a direct DNA association.

As circRNAs function through interacting with RBPs, these cacRNAs could also recruit RBPs that regulate transcription or splicing. Interestingly, CIRCpedia v3 reported the association of 120 and 100 proteins with *circDYNC1H1* and *circKIF2C,* respectively (**Supplementary Table S5**). To assess the relevant biological functions of these cacRNA-associated RBPs, we performed enrichment analysis on the genes of the RBPs. Interestingly, GO analysis of the RBPs associated with *circDYNC1H1* and *circKIF2C* indicated their involvement in several biological processes, including RNA processing, mRNA metabolic process, RNA biosynthetic process, mRNA processing, RNA splicing, and gene expression (**Supplementary Figures S8A and S8B)**. Furthermore, GO enrichment analysis of these RBPs for cellular component terms indicated that they are mainly localizing in ribonucleoprotein granule, nucleoplasm, nucleus, spliceosomal complex, and many others (**Supplementary Figure S9A and S9B)**. The GO molecular function terms for the RBP genes were RNA binding, mRNA binding, miRNA binding, mRNA 3’UTR binding, protein-RNA adaptor activity, N6-methyladenosine-containing RNA reader activity, among many others. **(Supplementary Figure S10A and S10B)**. Additionally, the KEGG pathway analysis revealed that the RBPs could be involved in several pathways like Spliceosome, mRNA surveillance pathway, Ribosome biogenesis in eukaryotes etc. **(Supplementary Figure S11A and S11B).**

To confirm the interaction between RBPs and cacRNA, we performed an RBP RNA immunoprecipitation (RIP) experiment. MBNL1 RIP experiment suggested an enrichment of *circDYNC1H1* and *circKIF2C* in MBNL1 pulldown samples, confirming the association of MBNL1 with these two circRNAs (**Figure 4H, I)**. Overall, the GO pathway analysis and pulldown experiment indicate that the RBPs associated with cacRNAs may regulate various nuclear events critical for gene expression.

## DISCUSSION

Since the last decade, the circRNA has become a hot topic among researchers due to its involvement in various diseases like cancer, autoimmune diseases, Alzheimer’s disease, cardiovascular disease, and diabetes [39, 40]. Although circRNA levels are lower than those of linear RNAs, they play a crucial role in cell proliferation, development, and differentiation by regulating gene expression at both transcriptional and posttranscriptional levels [40, 41]. CircRNAs execute their functions by interacting with different cellular regulators like genomic DNA, miRNAs, mRNAs, and proteins, or translate into polypeptides through cap-independent translation [6–9]. Although much has been studied about circRNA-mediated gene regulation via miRNAs or RBPs, the direct interaction of circRNAs with chromatin DNA is only now being discovered [18, 20]. The interaction of circRNA with chromatin and the regulation of gene expression remains largely unknown.

Various studies have demonstrated that RNAs interact with chromatin through either indirect interactions with proteins or direct sequence-specific interactions [18, 19, 42]. Although we identify hundreds of circRNAs associated with the chromatin fractions, their presence in this fraction due to their co-transcriptional biogenesis cannot be ruled out (**Figure 3**). To rule out the possibility of circRNA association with chromatin due to the co-transcriptional process, we terminated transcription with Actinomycin D treatment, followed by cellular fractionation. As expected, only a few cacRNAs were enriched in the chromatin fraction after transcription inhibition (**Figure 3**). Notably, the cacRNAs are also found in different cellular compartments, which may reflect the multimodal functions of the circRNAs. Moreover, two circRNAs-*circKIF2C* and *circDYNC1H1* were enriched in histone antibody pulldown, and *circDYNC1H1* was enriched in S9.6 antibody and RNaseH-GFP proteins that bind to DNA-RNA hybrids, indicating the direct interaction of these RNAs with chromatin DNA (**Figure 4**).

Given that, long stable R-loop formation is associated with harmful and stressful conditions, the presence of more circRNA-DNA hybrids could also reflect this [15]. Further research is required to identify the factors regulating circRNA-DNA hybrids and gene transcription at that locus across different pathophysiological conditions. In addition, our bioinformatics analysis suggested that the sequence complementarity between *circDYNC1H1* and different chromatin regions further confirms their chromatin interaction **(Figure 4)**. From this data, we can infer that circRNAs, by simultaneously interacting with multiple chromatin regions, may regulate the 3D chromatin architecture and gene expression. Even if the circRNAs may not play a direct role by themselves, the presence of associated RBPs with circRNAs may regulate several nuclear events in gene transcription and processing **(Supplementary Figures S8-S11)** [8]. In addition, found that in most cancers, the level of *circDYNC1H1* is generally lower than in normal tissue, and the *circKIF2C* level is highly upregulated in hepatocellular carcinoma (HCC), as reported by circAtlas3.0 **(Supplementary Figure 12)** [43]. The differential expression of these circRNAs across various cancers further underscores their functional role in maintaining cellular homeostasis. Collectively, these results suggest that circRNAs remained associated with cellular chromatin. Nonetheless, the overexpression or silencing of each cacRNA under physiological or pathological conditions would help elucidate the molecular mechanisms and functions.

In summary, our work identified and validated several chromatin-associated circRNAs. We also predicted and validated the RBPs associated with cacRNAs to execute their function. This finding warrants further research on cacRNA under pathophysiological conditions to understand its function. Overall, our study has opened a new avenue for investigating the function of circRNA and thereby further underscores its importance in gene regulation.

## DECLARATIONS

### Data availability

All the data generated in this study are included in the main text or supplementary data.

### Author contributions

Susovan Sadhukhan: Conceptualization, Methodology, Investigation, Formal analysis, Validation, Visualization, Writing-original draft, review, and editing. Komal Kumari: Methodology, Investigation, Formal analysis, Validation, Writing-review, and editing. Pranita Rout: Methodology, Investigation, Validation, Writing-review, and editing. Amaresh C. Panda: Conceptualization, funding acquisition, Supervision, Writing-original draft, review, and editing,

## Acknowledgements

We thank Dr. Punit Prasad for providing technical guidance in cellular fractionation. We thank our colleagues for their helpful discussions and manuscript proofreading. Some of the schematics were created with BioRender.com.

## Consent for publication

All authors consent this manuscript for publication.

