## Supplementary Figure for "Characterization of Chromatin-Associated Circular RNAs (cacRNAs)"

### SUPPLEMENTARY TABLE & FIGURES

**Supplementary table S1-** CacRNAs and nuclear circRNAs in HEK293 cells

**Supplementary table S2-** CacRNAs in H9 and Hela cells

**Supplementary table S3-** Oligonucleotide sequences used in this study

**Supplementary table S4-** CircRNA-genome BLAST data

**Supplementary table S5-** RBPs associated with cacRNAs

**Supplementary table S6-** GO terms and KEGG pathways of *circDYNC1H1* and *circKIF2C*-associated RBPs

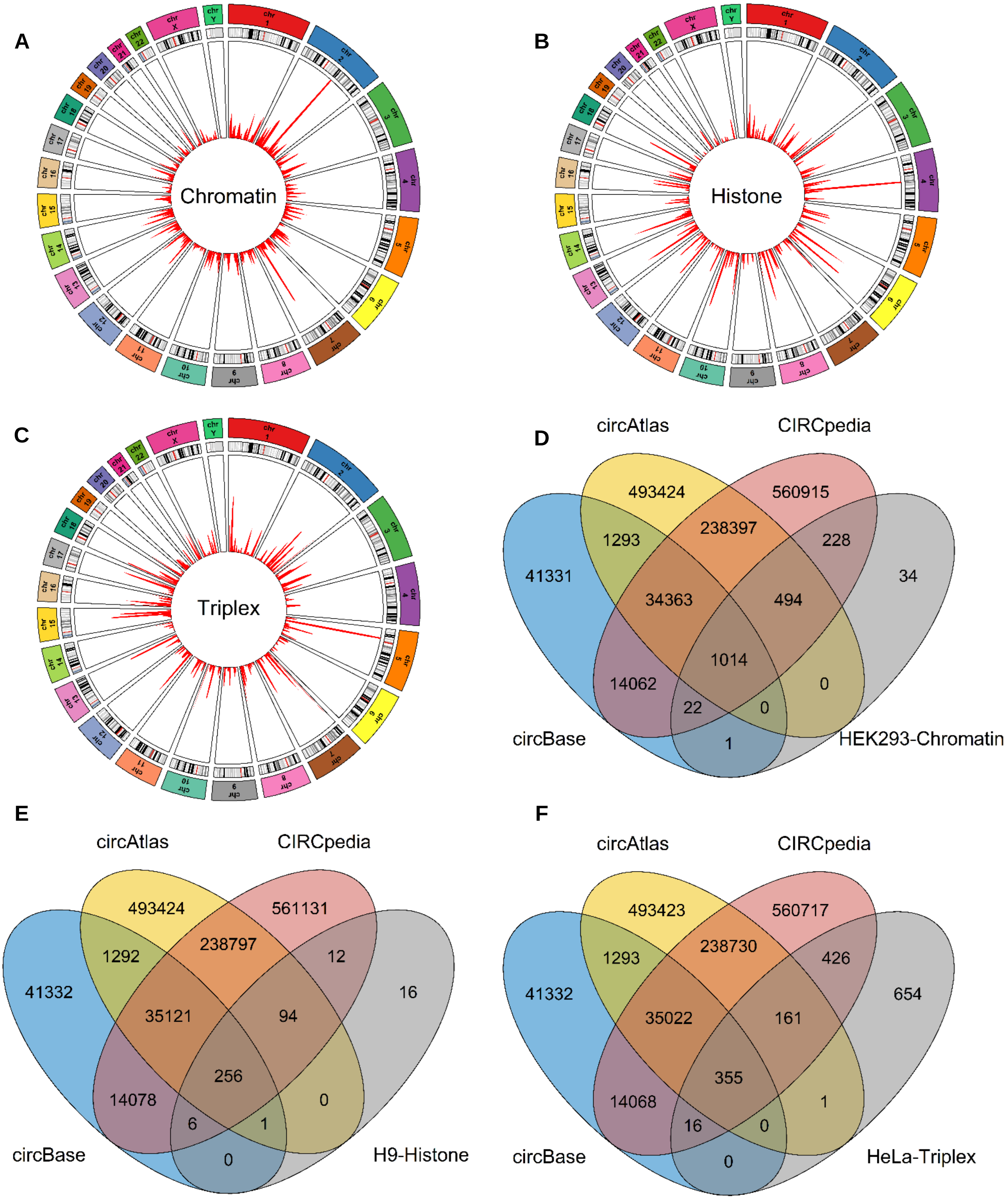


**Supplementary Figure S1: A.** Circos plot indicating the HEK293-Chromatin associated circRNAs originated from different human chromosomes. **B.** Circos plot indicating the H9-Histone associated circRNAs originated from different human chromosomes. **C.** Circos plot indicating the HeLa-DNA-RNA triplex forming circRNAs originated from different human chromosomes. **D-F.** Venn diagram between the circRNAs annotated in circBase, circAtlas3.0 and CIRCpedia v3 databases and the circRNAs analysed in HEK293-Chromatin (D), H9-Histone (E) and HeLa-DNA-RNA triplex (F) samples. The red arrows indicate the unique circRNAs we analysed in the respective samples.

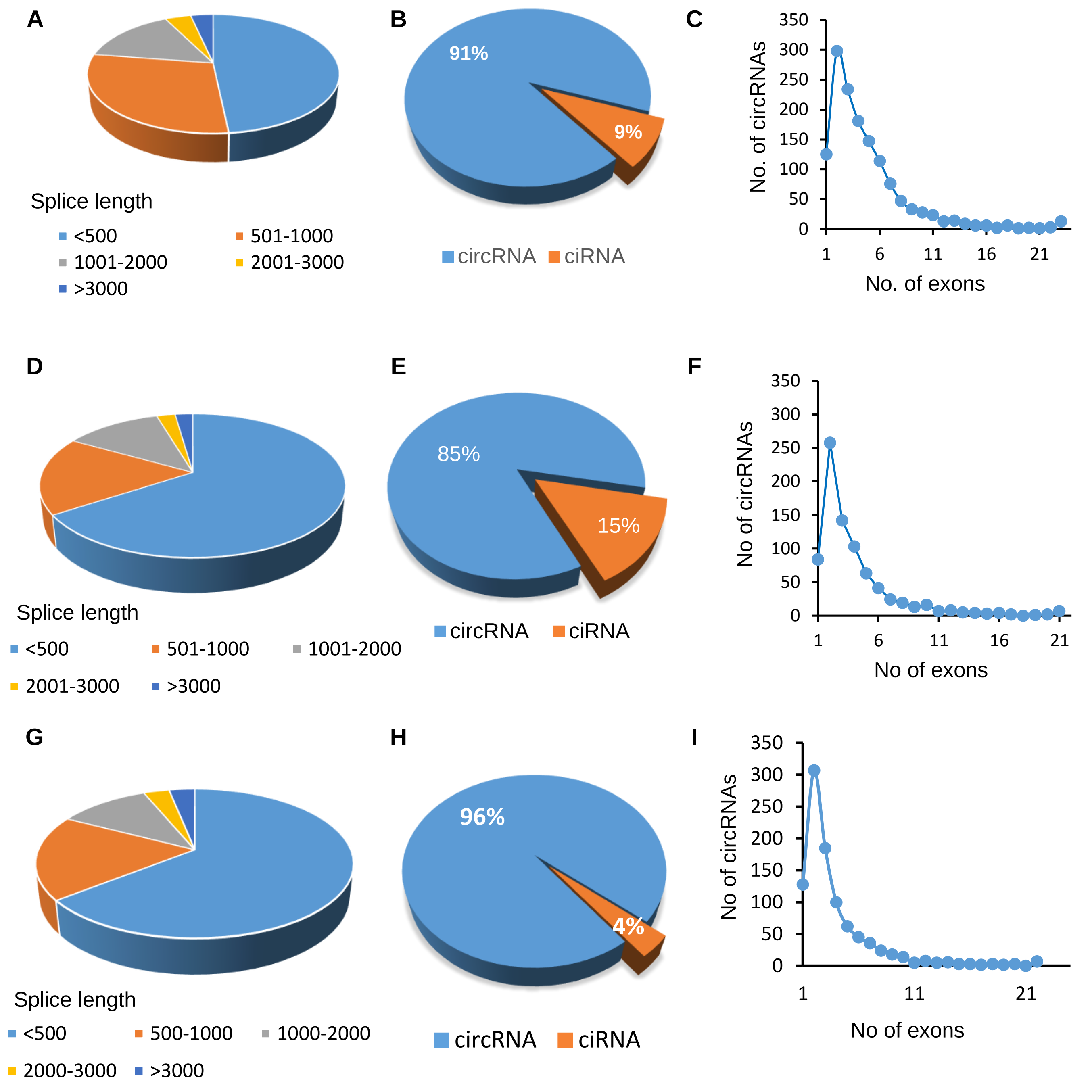


**Supplementary Figure S2: Characteristics of circRNAs identified in SNE, CPE and DRB samples.** **A.** Distribution of splice lengths of the circRNAs of SNE samples of HEK293 cells. **B.** Pie chart showing the number of exonic circRNAs and intronic ciRNAs identified in the SNE samples. **C.** Distribution of the exons in the exonic circRNAs of the SNE samples. **D.** Distribution of splice lengths of the circRNAs of CPE samples of HEK293 cells. **E.** Pie chart showing the number of exonic circRNAs and intronic ciRNAs identified in the CPE samples. **F.** Distribution of the exons in the exonic circRNAs of the CPE samples. **G.** Distribution of splice lengths of the circRNAs of DRB samples of HEK293 cells. **H.** Pie chart showing the number of exonic circRNAs and intronic ciRNAs identified in the DRB samples. **I.** Distribution of the exons in the exonic circRNAs of the DRB samples.

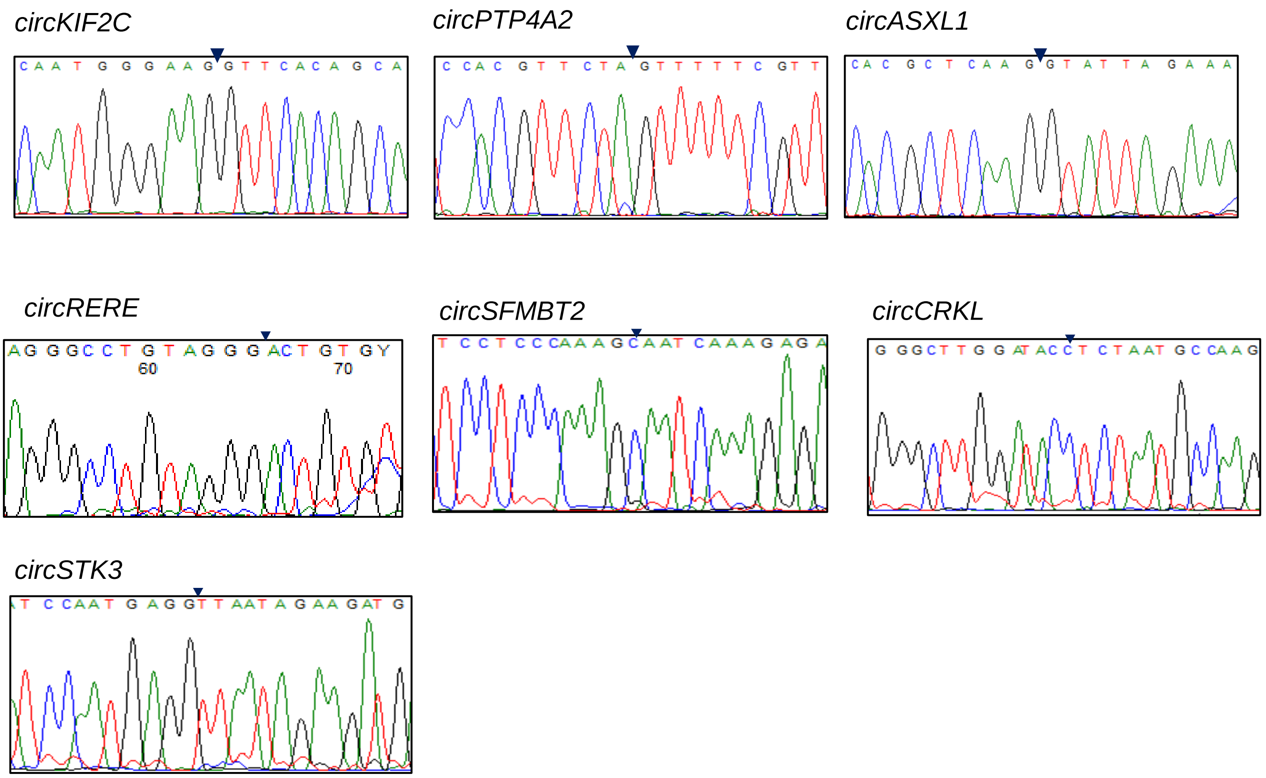


**Supplementary figure S3.** Sanger sequencing of the circRNA PCR products from HEK293T cells confirming their backsplice junction sequence, marked by arrow heads.

A

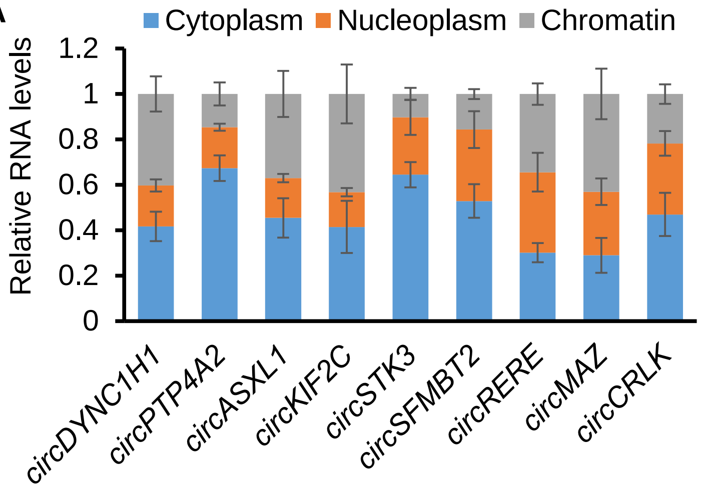


B

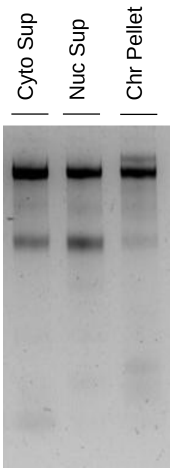


C

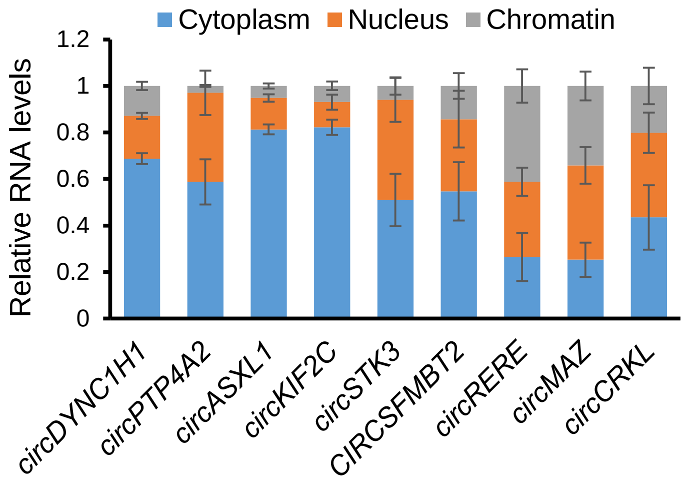


**Supplementary Figure S4: Abundance of circRNAs in different cellular fractions**. **A.** RT-qPCR analysis showing level of different circRNAs in the three cellular fractions in HEK293T cells. **B.** Total RNAs isolated from the three cell fractions- cytoplasmic supernatant, nuclear supernatant and chromatin pellet of Actinomycin D treated HEK293T cells, resolved on an agarose gel stained with SYBR-Gold. **C.** RT-qPCR analysis showing level of different circRNAs in the three cell fractions after ActD treatment of the HEK293T cells.

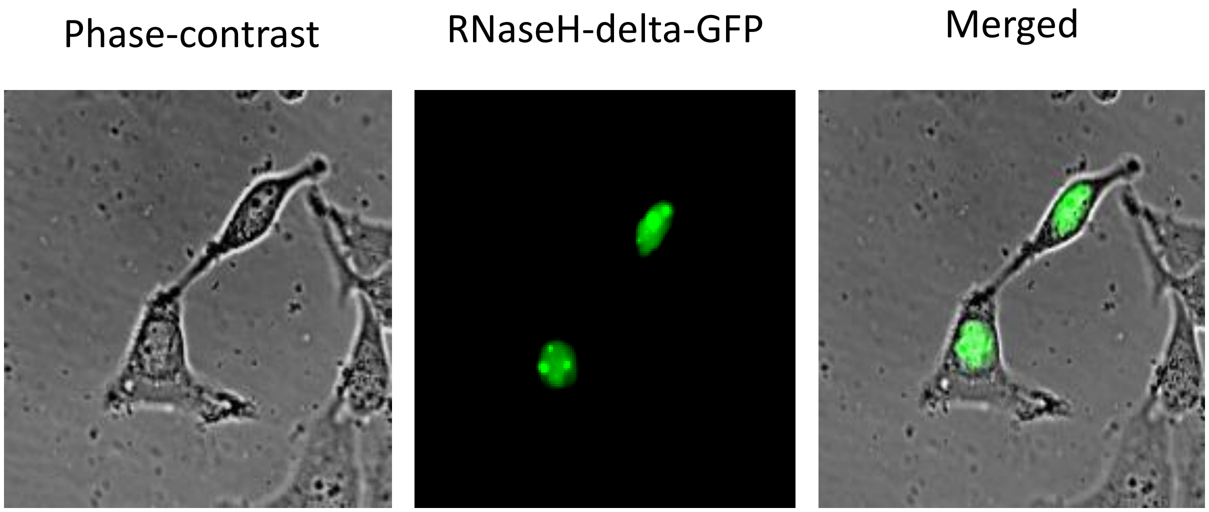


**Supplementary Figure S5:** Transfection of HEK293T cells with pEGFP-N2-2XNLS-RNaseH1 delta 1-27 (D210N) plasmid. The fluorescence imaging the expression of RNAseH-GFP in the cell nucleus.

**Sadhukhan et al, Supplementary Figure S6**

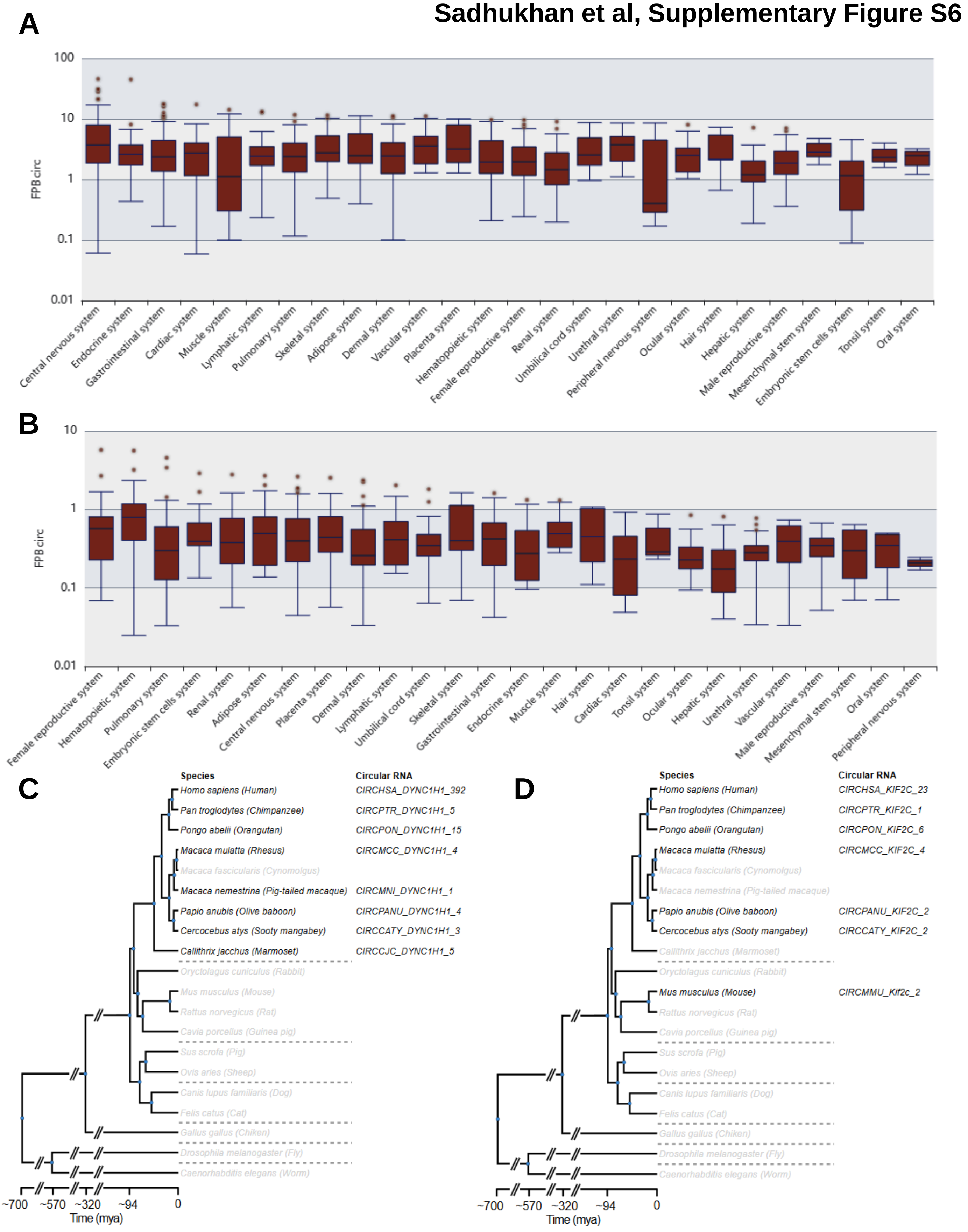


**Supplementary Figure S6: Expression and conservation analysis of *circDYNC1H1* and *circKIF2C* analyzed from CIRCpediav3**. **A.** The expression of *circDYNC1H1* in various tissue system of human. **B.** The expression of *circKIF2C* in various tissue system of human. **C.** The conservation of *circDYNC1H1*. **D.** The conservation of *circKIF2C*.

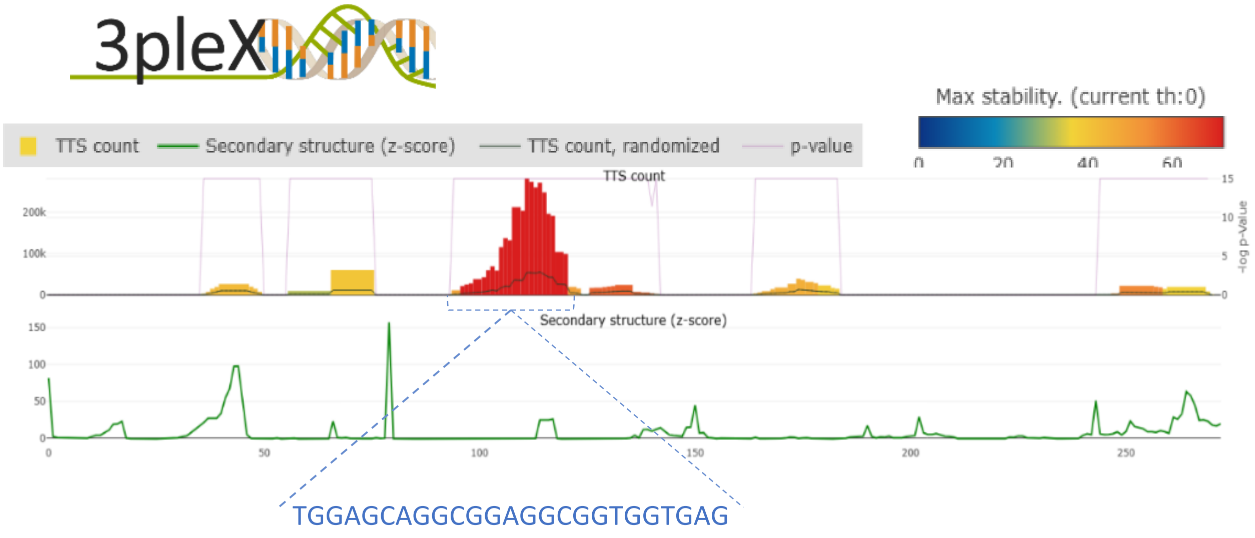


**Supplementary Figure S7: T**he triplex formation prediction of *circDYNC1H1* with human genomic sequences analyzed by 3plex predictor.

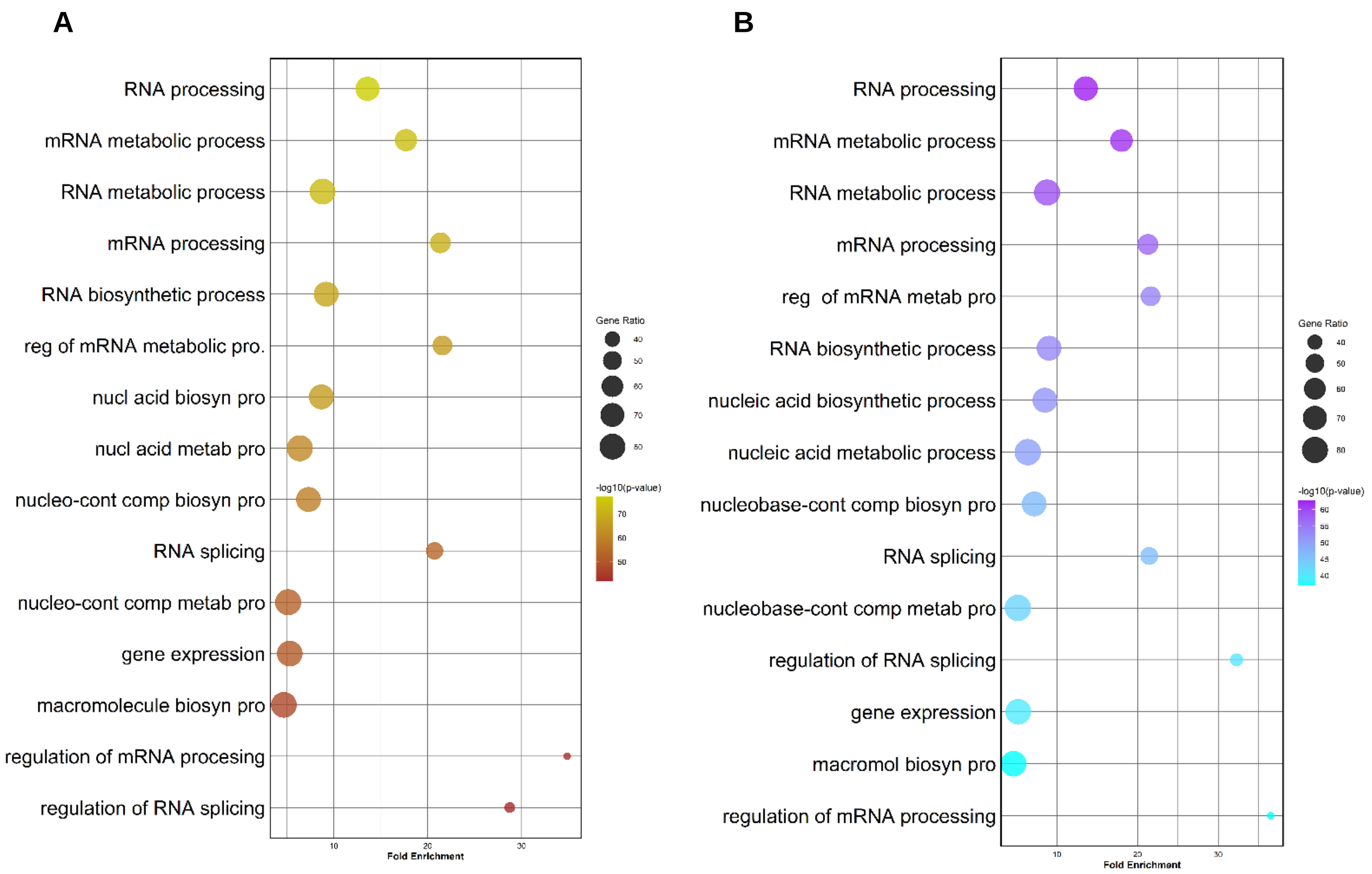


**Supplementary figure S8: Gene ontology (GO) Biological Process (BP) analysis**. Bubble plot showing the enriched GO terms for the Biological Process of the RBPs that are associated with *circDYNC1H1* (A) and *circKIF2C* (B).

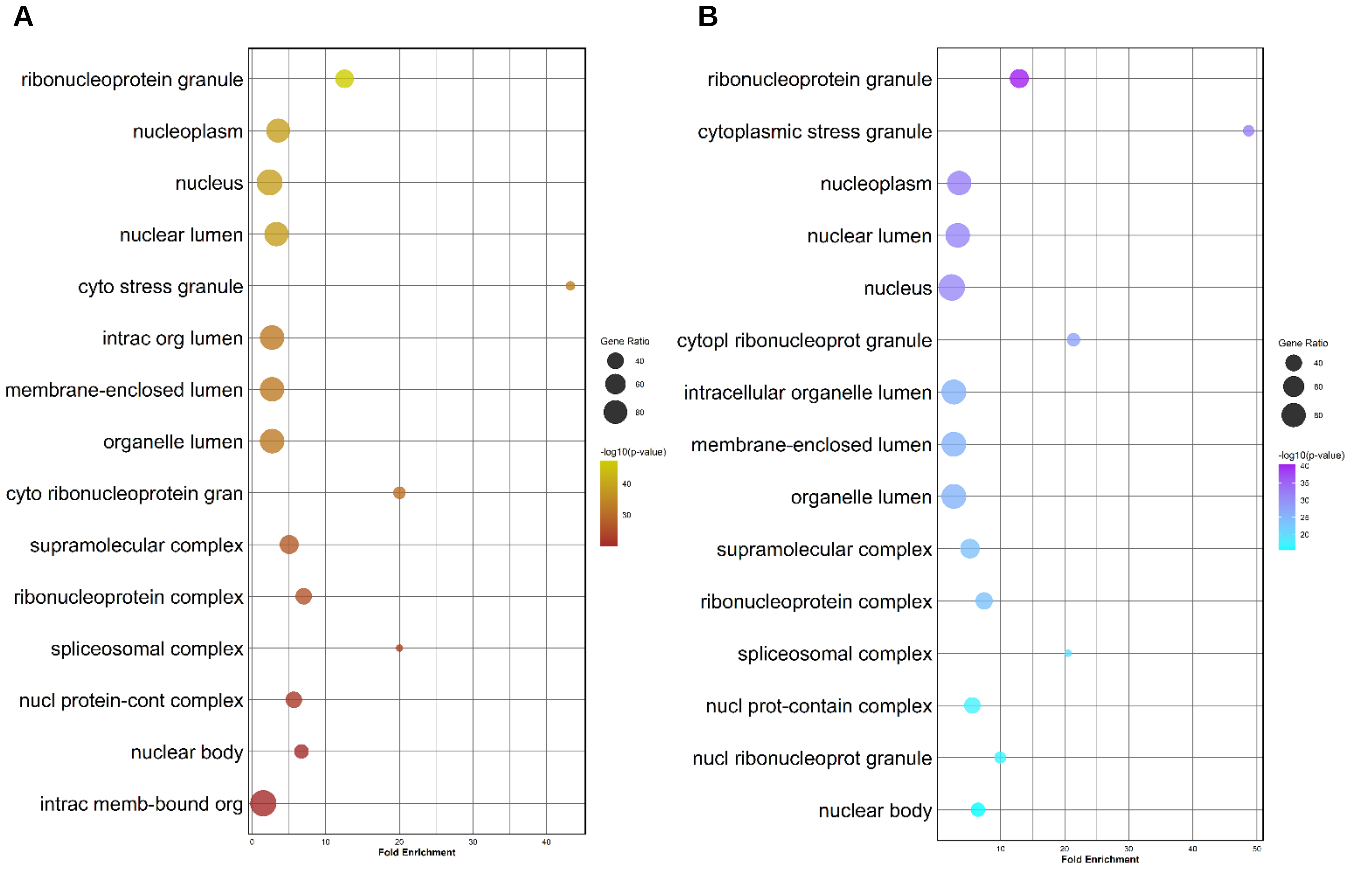


**Supplementary figure S9: Gene ontology (GO) Cellular component (CC) analysis**. Bubble plot showing the enriched GO terms for the Cellular component of the RBPs that are associated with *circDYNC1H1* (A) and *circKIF2C* (B).

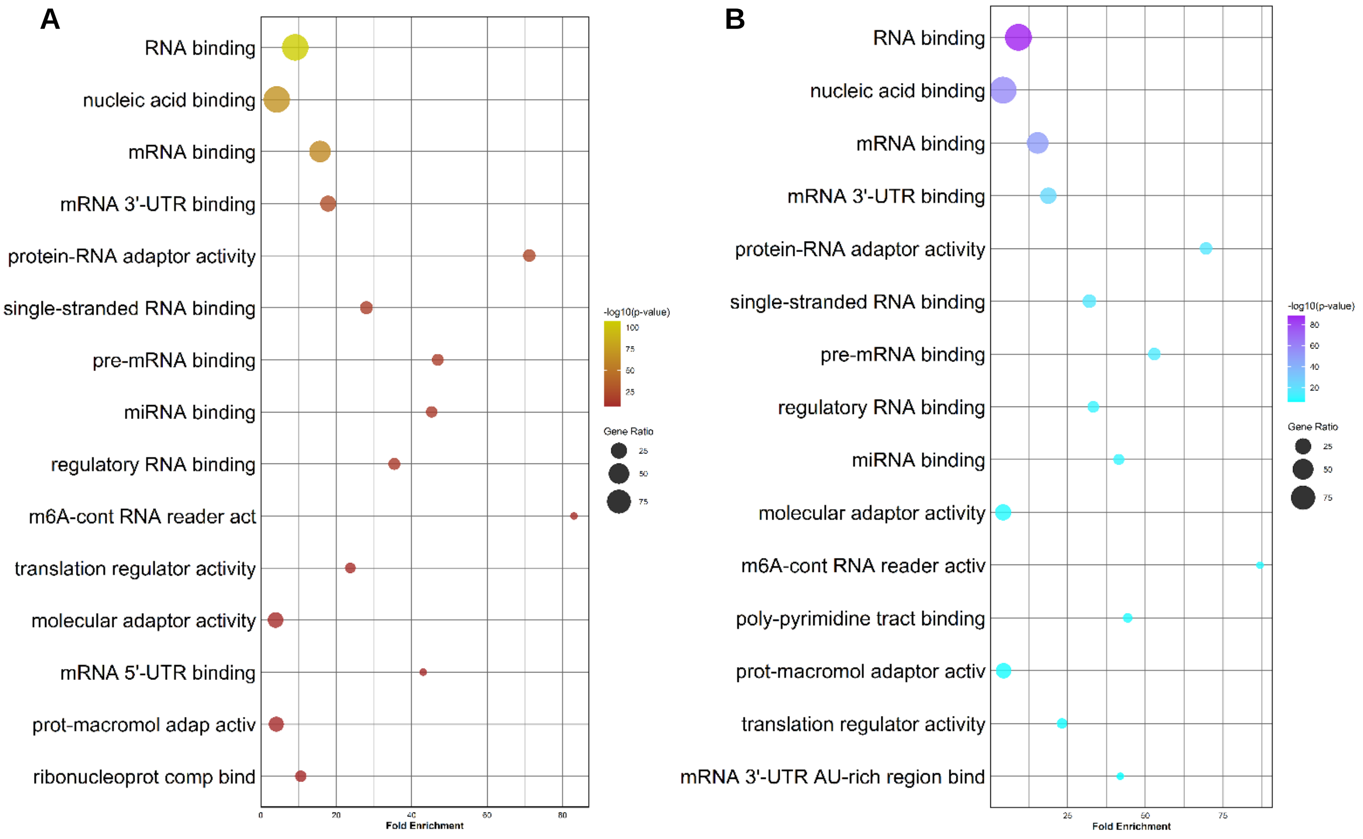


**Supplementary figure S10: Gene ontology (GO) Molecular functions (MF) analysis**. Bubble plot showing the enriched GO terms for the Molecular functions of the RBPs that are associated with *circDYNC1H1* (A) and *circKIF2C* (B).

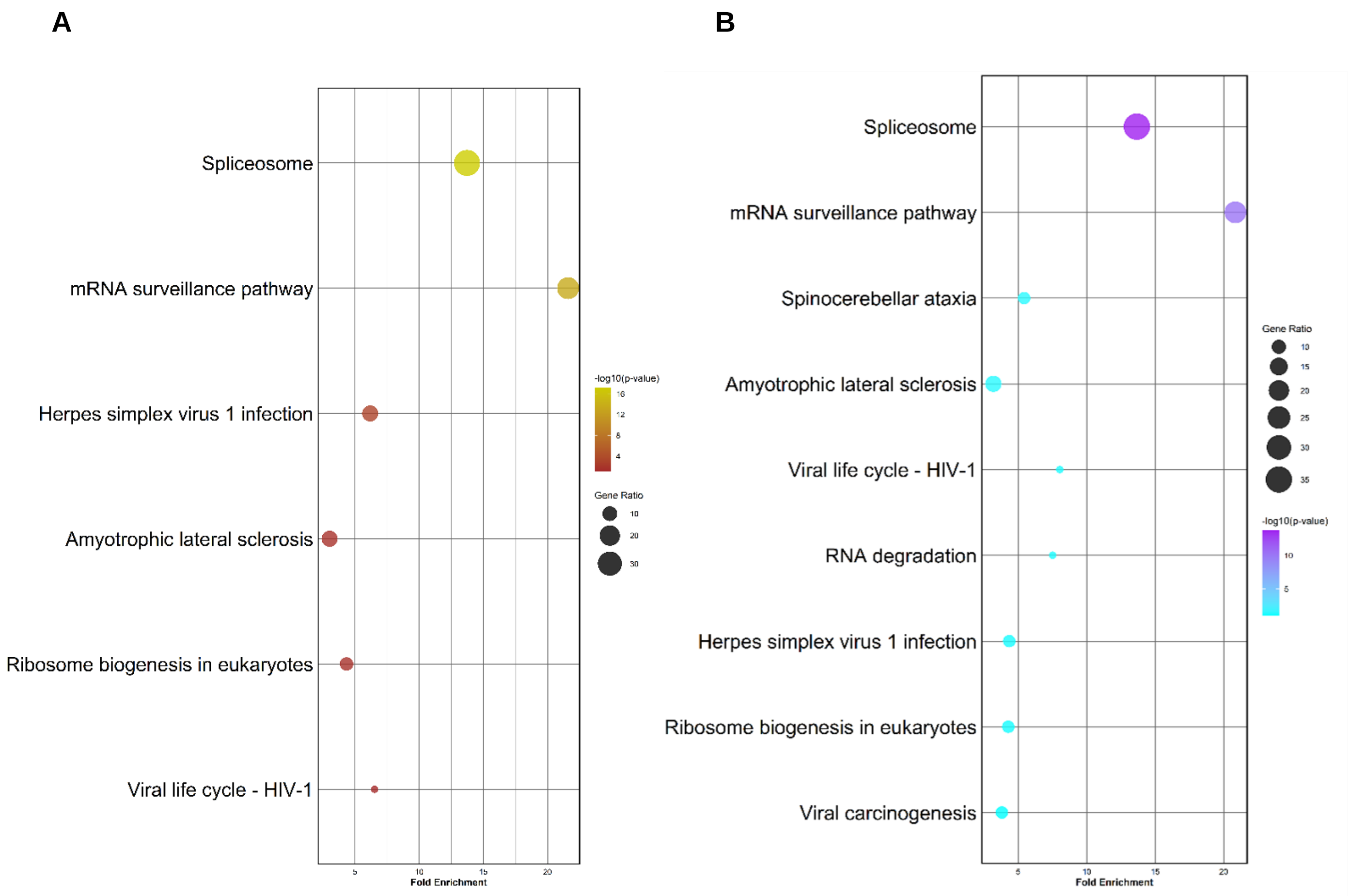


**Supplementary figure S11: KEGG pathway analysis**. Bubble plot showing the enriched KEGG pathways of the RBPs that are associated with *circDYNC1H1* (A) and *circKIF2C* (B).

A

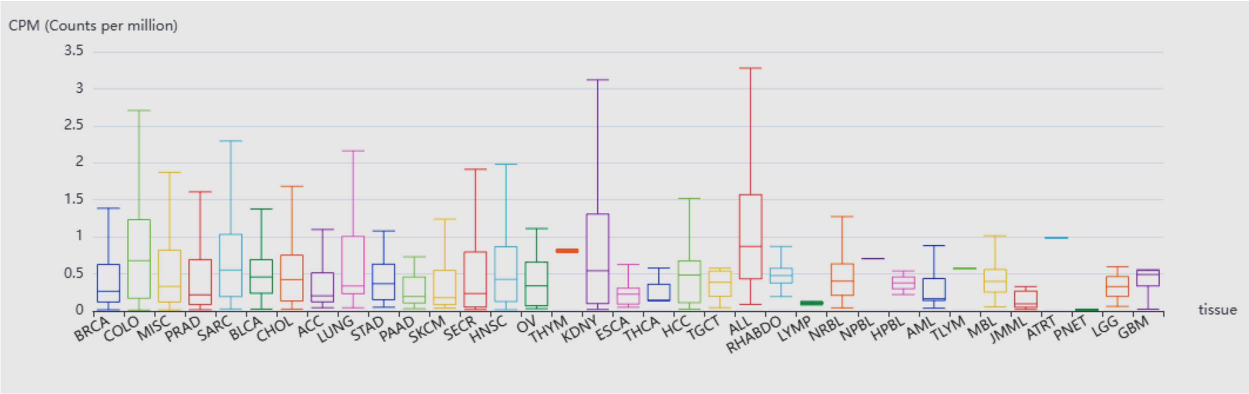


B

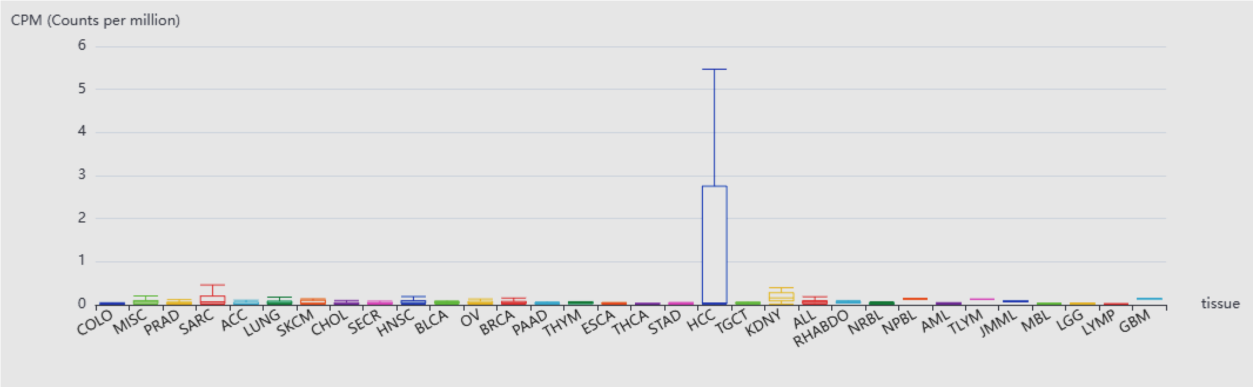


**Supplementary Figure S12: Expression of cacRNAs in cancers. A-B.** The expression of *circDYNC1H1* (**A**) & *circKIF2C* (**B**) expression in various cancers reported by circAtlas3.0.
